# An Analog-Sensitive Tau Tubulin Kinase 2 Enables Acute Chemical-Genetic Control of Primary Cilia

**DOI:** 10.64898/2026.10.09.757601

**Authors:** Andrea Lacigová, Lukáš Čajánek

## Abstract

Kinases regulate a broad spectrum of cellular processes, including the formation of primary cilia, microtubule-based organelles that protrude from the cell surface and play important roles in development and disease. TTBK2 is a key ciliary kinase that controls primary cilium assembly and maintenance. However, highly selective small-molecule inhibitors of TTBK2 are still lacking, representing a major bottleneck for investigating the temporal requirements and dynamics of TTBK2-dependent processes in cilia.

Here, we report the first analog-sensitive variant of TTBK2, enabling rapid, reversible, and highly specific control of TTBK2 kinase activity in ciliogenesis-relevant hTERT-RPE1 cells. Using this system, we define the kinetics of CEP164 phosphorylation, cilium shortening and reassembly, and IFT88 retention at the ciliary base following acute TTBK2 inhibition and subsequent inhibitor washout.

Together, our findings provide new insight into TTBK2-dependent pathways and establish analog-sensitive TTBK2 as a powerful tool for investigating the dynamics of TTBK2-regulated processes with high temporal resolution.

## Introduction

The primary cilium is a hair-like structure on the surface of most vertebrate cells. In contrast to motile cilia or flagella, primary cilia are immotile. Despite their lack of motility, they are responsible for conveying a wide range of extracellular signals to downstream intracellular signaling pathways, making them crucial regulators of both embryonic development and tissue homeostasis in adulthood (Čajánek et al. 2025; Hilgendorf et al. 2024; Mill et al. 2023). While severe, systemic disruption of ciliary signaling is considered embryonic lethal, a broad spectrum of defects in ciliary assembly or function underlie a diverse group of human diseases collectively known as ciliopathies (Hildebrandt et al. 2011; Wallingford 2019). A cilium is composed of the basal body, transition zone, and a microtubule-based axoneme surrounded by the ciliary membrane. The basal body is derived from the mother centriole, the older of the two centrioles that comprise the centrosome. In turn, centrioles are barrel-shaped organelles exhibiting a characteristic nine-fold radial symmetry (Breslow and Holland 2019).

Tau Tubulin Kinase 2 (TTBK2), a distant member of the CK1 kinase family, plays an instrumental role through its kinase activity in both the initiation of ciliogenesis and the maintenance of assembled cilia at the appropriate length (Goetz et al. 2012; Lacigová and Čajánek 2025; Nguyen and Goetz 2022). Specifically, TTBK2 regulates the formation of the ciliary vesicle at the distal end of the future basal body/mother centriole (Benk Vysloužil et al. 2025; Kanie et al. 2025; Lo et al. 2019), which is considered the earliest event in the cilium assembly pathway (Sorokin 1962). In addition, TTBK2 activity mediates the loss of protein CP110 from the mother centriole (Goetz et al. 2012). The role of CP110 is to inhibit microtubule growth at the centriole distal end (Iyer et al. 2025). In turn, its removal from the mother centriole represents another hallmark of the initiation of ciliogenesis and is considered a prerequisite for axoneme extension (Spektor et al. 2007). The growth and maintenance of the ciliary axoneme are supported by the activity of Intraflagellar Transport (IFT), a process driven by IFT protein complexes in coordination with kinesin and dynein motors that shuttle cargo between the ciliary tip and base (Lacey and Pigino 2025). Loss of TTBK2 impairs association of IFT proteins with the mother centriole distal end, hence effectively disrupting IFT (Benk Vysloužil et al. 2025; Čajánek and Nigg 2014; Goetz et al. 2012).

One of the unresolved questions regarding TTBK2 function during ciliogenesis concerns the timing and dynamics of the individual events regulated by its kinase activity. Although small molecules capable of inhibiting TTBK2 have been described (Bashore et al. 2023; Halkina et al. 2021; Flax et al. 2026), their notable drawback is the inability to selectively target TTBK2 over TTBK1 owing to the high similarity of their kinase domains. Given that TTBK1 activity has also been proposed to contribute to ciliogenesis in specific contexts (Binó and Čajánek 2023), alternative means of acutely inhibiting TTBK2 kinase activity are required. Analog-sensitive kinase technology is a chemical-genetic strategy that enables selective inhibition of individual kinases by bulky ATP analogs (Bishop et al. 2000; Lopez et al. 2014). It is based on mutation of a structurally conserved amino acid within the kinase active site, termed the gatekeeper residue, to a smaller residue. The resulting enlargement of the ATP-binding pocket creates a unique binding site not present in wild-type kinases. Consequently, the engineered analog-sensitive kinase can be specifically inhibited by bulky ATP-competitive inhibitors, such as PP1 derivatives bearing a C3 substituent, which fit the enlarged ATP-binding pocket but are unable to bind wild-type kinases with larger gatekeeper residues, such as methionine or phenylalanine. This strategy has been successfully applied to dozens of kinases (Blethrow et al. 2004; Hanke et al. 1996; Lopez et al. 2014).

In this study, we report adopting the analog-sensitive strategy to develop a model that enables specific inhibition of TTBK2 in ciliogenesis-relevant hTERT-RPE1 cells. To our knowledge, this represents the first example of an engineered ATP analog-sensitive kinase from the CK1 family. This model should prove valuable for investigating the dynamics of TTBK2-regulated events with high temporal resolution and for elucidating their underlying molecular mechanisms.

## Materials and Methods

### Cell culture, stable cell line generation, and treatments

HEK293T cells were cultivated in DMEM cultivation media (ThermoFisher, Cat.N. 31966047) supplemented with 10% fetal bovine serum (BioTech, Cat.N. FB-1101/100) and 1x penicillin/streptomycin (BioTech, Cat.N. XC-A4122) (DMEM complete media). hTERT-RPE1 cells were cultivated in DMEM/F-12 cultivation media (ThermoFisher, Cat.N. 21331020) supplemented with 10% fetal bovine serum (BioTech, Cat.N. FB-1101/100), 1x penicillin/streptomycin (BioTech, Cat.N. XC-A4122), and 2 mM L-glutamine (BioTech, Cat.N. XC-T1715) (DMEM/F-12 complete media). To induce expression of genes under the doxycycline (DOX)-inducible promoter, DOX (MedChem, Cat.N. HY-N0565B) was added to the culture media to a final c=200 nM, unless stated otherwise.

To generate stable cell lines, TTBK2 KO hTERT-RPE1 (Bernatik et al. 2020) cells were seeded on 6 cm Petri dishes to be ∼60 % confluent on day 2. On day 2, cells were transfected with 500 ng of pgLAP2 NEO plasmid (Binó et al. 2022) containing TTBK2 M94A or TTBK2 M94G, 5.5 µg of pOG44 plasmid, and DOX was added to the media. For transfection, Lipofectamine 3000 (ThermoFisher, Cat.N. L3000001) kit was used according to the manufacturer’s instructions. 24 hours after transfection, media was changed and G418 antibiotic (in final c=350 μg/ml, Merck, Cat.N. G8168) was added. The media containing DOX and G418 was changed every 48 hours for a period of 2 weeks post transfection to select clones with stably integrated genes of interest. Selected colonies were further cultivated and validated.

To validate the expression of Flag-TTBK2 M94A or Flag-TTBK2 M94G, 100 000 cells were seeded per well in a 24-well plate in tetracycline-free media with or without DOX, cultivated for 48 hours, and then lysed and analyzed by Western blot (WB).

To evaluate the rescue of cilia formation in these cell lines, 100 000 cells were seeded on coverslips in 24-well plates and cultivated for 72 hours in media with 10 % FBS, followed by 24 hours of cultivation in media without FBS, and the cells were then fixed and stained. DOX was added to the culture media throughout the entire duration of the experiment.

To initially evaluate inhibition of TTBK2 M94G kinase activity, TTBK2 KO HEK293T cells (Bernatik et al. 2020) were seeded on a 24-well plate (100 000 cells/well). After 24 hours, the cells were transfected using PEI (3.5 μl of 2 mg/ml stock solution of PEI was used for 1 μg of plasmid DNA) with 100 ng of DVL2-MYC plasmid and 400 ng of either Flag-TTBK2 WT, Flag-TTBK2 M94G, or pcDNA3.1 plasmid, and DOX was added. 24 hours post transfection, cells were treated with the respective ATP-analog compound in indicated concentrations or with DMSO. After 24-hour treatment, cells were lysed and analyzed on WB.

Throughout the experiments, the following ATP analogs were used: 1-NM-PP1 (PubChem CID: 5154691, Merck, Cat.N. 529581), 3-MB-PP1 (PubChem CID: 24865390, Merck, Cat.N. 529582), and 3-IB-PP1 (PubChem CID: 25243798, Merck, Cat.N. 529598), with DMSO treatment used as a negative control.

### Cloning and mutagenesis

pENTR-TOPO-TTBK2 Gateway entry plasmid (Bernatik et al. 2020) was used as a template to introduce M94A (forward primer: cagattccgaccctgcaactgggcgaccacatagttgaatcgatc, reverse primer: gatcgattcaactatgtggtcgcccagttgcagggtcggaatctg) and M94G (forward primer: ttccgaccctgcaactgcccgaccacatagttgaatcg, reverse primer: cgattcaactatgtggtcgggcagttgcagggtcggaa) substitutions using QUIKChange II XL side-directed mutagenesis kit (Agilent Technologies, Cat.N. 200522). PCR products were transformed into XL10-Gold chemically competent bacteria (Agilent Technologies, Cat. no. 200315), and the introduced mutations were verified by DNA sequencing. The inserts were cloned into pgLAP2 NEO Gateway destination vector using LR Clonase II Enzyme mix (ThermoFisher, Cat. N.11791020).

### Inhibition of ciliogenesis and inhibitor wash-out experiments

For complete inhibition of ciliogenesis, 80 000 cells were seeded on coverslips in 24-well plates into media containing DOX. On day 2, ATP-analog/DMSO was added to the media as indicated. On day 3, media was changed to low-serum (1% FBS) media and appropriate ATP-analog/DMSO treatment was added. On day 4, cells were fixed and stained.

For inhibitor wash-out experiments evaluating ciliogenesis (Fig.3B-F, Suppl. Fig.3B-D), 80 000 cells were seeded on coverslips in 24-well plates into media containing DOX. On day 2, 3-MB-PP1/DMSO was added to the media as indicated (10 μM concentration in Suppl. Fig.3C-D, 25 μM concentration in Fig.3B-F). On day 3, media was changed to low-serum (1% FBS) media and appropriate 3-MB-PP1/DMSO treatment was added. On day 4, media was changed, and ATP-analog was washed out from the cells for various periods of time as indicated, and these cells were treated with DMSO until fixation (Fig.3D-F). All conditions were then fixed and stained. In the case of Fig.3B-C and Suppl. Fig.3B-D, cells were cultivated for another 24 hours and were fixed on day 5.

### Cilia maintenance experiments

For experiments depicted in Suppl. Fig.4B-D, 80 000 cells were seeded on coverslips in 24-well plates into media containing DOX. On day 3, media was changed to low-serum (1% FBS) media. On day 4, cells were either fixed or further cultivated for 24 hours in low-serum media with addition of 3-MB-PP1 (25 μM) or DMSO. On day 5, the cells were fixed and stained.

For experiments depicted in Fig.4 and Suppl. Fig.4E-H, 80 000 cells were seeded on coverslips in 24-well plates into media containing DOX. On day 2, media was changed to low-serum (1 % FBS) media. On day 3, cells were treated with 3-MB-PP1 (25 μM) or DMSO as indicated and then fixed and stained.

### CEP164 phosphorylation

80 000 cells were seeded in a 24-well plate into media containing DOX. For the experiment depicted in Suppl. Fig.3E, 3-MB-PP1 (25 μM) or DMSO was added to the cells for various periods of time as indicated on day3 and the cells were then lysed and analysed by WB. For the experiment depicted in Fig.3G, 3-MB-PP1 (25 μM) or DMSO was added to the cells on day 2, and on day 3, the treatment was washed out from the cells for various periods of time as indicated. The cells were then lysed and analysed by WB.

### Western blot

Cells grown in 24-well plates were lysed in 100 μl of 1x Laemmli buffer (31.5 mM Tris-HCl (pH 6.8), 2% SDS, 10% glycerol, 5% β-mercaptoethanol, 0.01% bromophenol blue). Collected lysates were denatured at 95 °C for 15 min, loaded onto a 5% acrylamide stacking gel combined with an 8% running gel, and ran for 80 min at 150 V in running buffer (0.1% SDS, 0.192M glycine, 0.025M Tris-base). The proteins were then transferred onto an Immobilon-PVDF membrane (Millipore, Cat.N. IPVH00010) for 85 minutes at 100 V in transfer buffer (20% methanol, 0.192 M glycine, 0.025 M Tris-base). Membranes were then blocked for 1 hour in 5% skimmed milk in wash buffer (20mM Tris-base, 0.1% Tween 20, 150mM NaCl) followed by overnight incubation at 4 °C with the following primary antibodies diluted in 5% skimmed milk: CEP164 (22227-1-AP/Proteintech) 1:500; DVL2 (3216/Cell Signaling) 1:500; TTBK2 (HPA018113/Sigma) 1:500; α-tubulin (11224-1-AP/Proteintech)1:1000). The membranes were then washed 3x for 10 minutes in wash buffer on a shaker and then incubated for 2 hours at RT with secondary antibodies (HRP-linked anti-rabbit IgG (7074S/Cell Signaling) 1:1000-1:500) diluted in 5% skimmed milk and washed 3x for 10 minutes in wash buffer. The membranes were then developed using ECL Prime (Cytiva, Cat.N. RPN2236) and ChemiDoc Imaging System (Bio-Rad, Cat.N.12003154). For every membrane, a labelled protein ladder (Thermo Fisher, Cat.N.26625) was used.

### Immunofluorescence microscopy and image analysis

For immunofluorescent analysis, cells grown on coverslips on a 24-well plate were washed with PBS and fixed using ice-cold methanol at -20°C for 10 minutes. The coverslips were then briefly washed with PBS, incubated for 1 hour at RT in blocking buffer (2% BSA, 0.02% NaN_3_ in 1x PBS), then incubated for 1 hour at RT with primary antibodies (ARL13B (17711-1-AP/Proteintech) 1:1000; CEP164 (22227-1-AP/Proteintech) 1:500; CP110 (140-195-5/Sigma)1:25; IFT88 (13967-1-AP/Proteintech) 1:500; TTBK2 (HPA018113/Sigma) 1:500; γ-tubulin (T6557/Sigma) 1:1000) diluted in blocking buffer, washed 3x for 10 minutes with PBS before being incubated for 1 hour at RT in a dark chamber with secondary antibodies and Hoechst dye diluted in blocking buffer (Alexa488 anti-rabbit (A21206/ThermoFisher) 1:1000; Alexa568 anti-mouse (A10037/ThermoFisher) 1:1000; Hoechst (62249/ThermoFisher) 1:2000), washed 3x10 minutes with PBS again and mounted using ProLong Glass Antifade Mountant (ThermoFisher, Cat.N. P36980). In case of triple staining (Fig.2, Fig.3, Suppl. Fig.2, Suppl. Fig.3), the cells were incubated for 1 hour at RT with CETN1 antibody conjugated with Alexa647 (12794-1-AP/Proteintech) dye prior to the final wash steps and mounting. The images were acquired with the use of ZEISS AxioImager.Z2 microscope equipped with Plan-Apochromat 100x/1.40 Oil DIC objective and a monochromatic Hamamatsu ORCA Fusion camera. Raw images were acquired as z-stacks and further processed using ImageJ software (Schneider et al. 2012).

Analysis of the microscopic images was performed on maximum intensity projections from the acquired z-stacks. ARL13B-stained cilia were counted and measured manually using the segmented line tool. For quantification of ciliated cells, the identified cilia per image were divided by the number of centrosomes and multiplied by 100. Signal intensities of CP110 were measured from an area (circle with 0.58 μm diameter) around the CEP164-positive centriole, and background intensity (from the same-size area within the cell, but outside of the centrosome) was subtracted. Signal intensities of IFT88 were measured from an area (circle with 0.87 μm diameter) around the centrosome (with subtracted background intensity from the same-size area within the cell) and normalized by the signal intensity (with subtracted background intensity) of γ-tubulin staining within the same area.

### Statistical analysis and software

The statistical analyses (one-way ANOVA with Tukey’s multiple comparisons, or unpaired Student’s T-test) and data visualization were performed with GraphPad Prism 8 software. The data are presented as mean ±SD unless stated otherwise. Individual data points represent either field of view (in case of quantification of ciliated cells) or individual cilia/centrosomes.

The AlphaFold modelling was done using freely available AlphaFold server (https://alphafoldserver.com). The models were run with default settings. The protein sequence for modelling of TTBK2 kinase domain (residues 21-284) is available in Uniprot database (https://www.uniprot.org) with the analog-sensitive variants (M94A and M94G) possessing a single M at position 94 changed to A or G, respectively. Visualisation of the models was performed in the ChimeraX software, with surface displayed for the whole model and highlighted the residue at position 94 (M in WT or G in mutant variant).

The graphics in this paper were created using Adobe Illustrator 2021 software.

## Results and Discussion

### Generation of ATP analog-sensitive variant of TTBK2

To engineer an ATP-analog variant (Figure 1A) of TTBK2, we inspected its kinase domain. Following the published guidelines (Blethrow et al. 2004) and using AlphaFold modeling, we identified the methionine at position 94 as the so-called gatekeeper residue residing in the hydrophobic pocket of the kinase domain (Figure 1B). Generally, the bulky gatekeeper residue can be substituted with either glycine or alanine to alter the shape of the pocket and induce sensitivity to ATP analogs such as derivatives of PP1. We therefore generated both alanine- and glycine-variants of analog-sensitive TTBK2, here termed TTBK2 M94A and TTBK2 M94G (Figure 1B).

**Figure 1:**
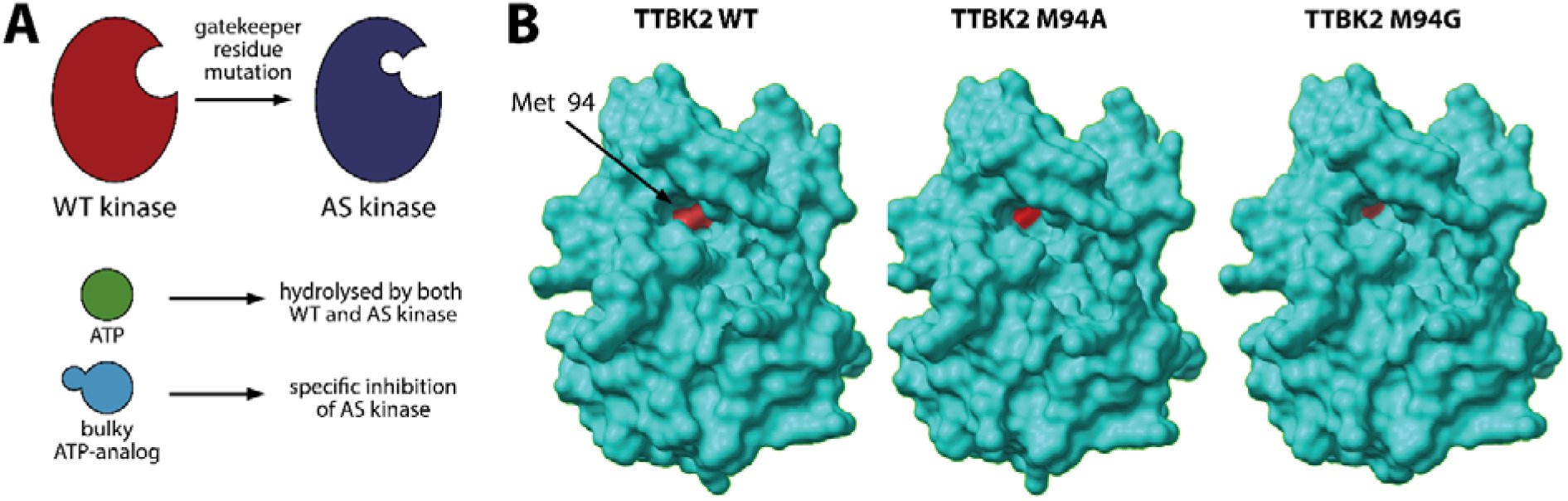
**(A)** Principle of ATP-analog sensitive kinase technology. Gatekeeper residue mutation in a given kinase domain results in sensitivity to inhibition by a synthetic non-hydrolysable ATP-analog while preserving catalytic activity in its absence. **(B)** AlphaFold models of the TTBK2 kinase domain, including the methionine at position 94 (M94) identified as gatekeeper. TTBK2 WT and mutants where the gatekeeper was replaced by alanine (M94A) or glycine (M94G) are shown. The gatekeeper residue or its substituents are highlighted in red.

Next, we examined three commercially available ATP-analog compounds, 1-NM-PP1, 3-MB-PP1, and 3-IB-PP1 in their ability to inhibit the ATP-analog-sensitive TTBK2. At this point, we focused on TTBK2 M94G, as glycine substitutions of the gatekeeper residue typically demonstrate higher sensitivity to ATP analogs over alanine-based substitutions (Blethrow et al. 2004; Lopez et al. 2014). We co-transfected TTBK2 KO HEK293 cells (Bernatik et al. 2020) with Flag-TTBK2 WT or M94G, together with DVL2, a WNT pathway component undergoing phosphorylation and mobility shift in the presence of TTBK2 (Bernatik et al. 2020; Hanáková et al. 2019). We observed that both TTBK2 WT and M94G induced a mobility shift in DVL2 (Figure S1A), suggesting that the catalytic activity of TTBK2 M94G is not dramatically impaired by the gatekeeper mutation. Moreover, we observed an effect of 3-IB-PP1 on DVL2 mobility in 50 and 10 μM concentrations, and an effect of 3-MB-PP1 in 50 μM concentration. Treatment with 1-NM-PP1 showed no apparent effect in any concentration tested, indicating a limited potency of this compound to inhibit TTBK2 M94G. In turn, 1-NM-PP1 was excluded from all further experiments.

Kinase activity of TTBK2 is essential for primary cilia formation (Bernatik et al. 2020; Goetz et al. 2012). As our results indicated an inhibitory effect of 3-MB-PP1 and 3-IB-PP1 on the activity of ATP-analog-sensitive TTBK2 (M94G variant), we wanted to corroborate this finding using a ciliogenesis-related readout. To this end, we used hTERT-RPE1 TTBK2 KO cells (Bernatik et al. 2020) to establish DOX-inducible cell lines expressing Flag-TTBK2 M94G or Flag-TTBK2 M94A. We examined two clonal cell lines for both TTBK2 M94G and TTBK2 M94A, respectively. Having confirmed that these cell lines express the respective TTBK2 variants in the presence of DOX (Figure S1B), we assessed the ability of TTBK2 M94G and M94A to rescue cilia formation in hTERT-RPE1 TTBK2 KO cells following serum starvation (Figure S1C). Importantly, all the tested clones showed comparable ciliation rates as well as the length of assembled primary cilia (Figure S1D-F), demonstrating that both M94G and M94A TTBK2 mutants possess sufficient catalytic activity to support ciliogenesis. Gatekeeper substitutions by glycine exhibit higher sensitivity to ATP analog treatment, yet sometimes at the expense of overall kinase activity (Lopez et al. 2014; Blethrow et al. 2004). As the latter did not seem to be the case for ciliogenesis-related activity of TTBK2, we focused on the TTBK2 M94G variant from now on.

To investigate the consequences of pharmacological inhibition of TTBK2 M94G for primary cilia formation, we treated the cells with either 3-MB-PP1 or 3-IB-PP1 for 48 hours (10 or 25 μM dose), and serum-starved them for the last 24 hours (Figure 2A). hTERT-RPE1 TTBK2 KO cells, inducibly expressing Flag-TTBK2 WT (Bernatik et al. 2020) were used as a control. We observed that both 3-MB-PP1 and 3-IB-PP1 treatment significantly reduced the number of ciliated cells, if compared to either DMSO-treated TTBK2 M94G cells or to 3-MB-PP1/3-IB-PP1-treated TTBK2 WT condition (Figure 2B and 2D). Importantly, neither 3-MB-PP1 nor 3-IB-PP1 treatment affected the ciliation rate in control cells expressing TTBK2 WT, demonstrating the effect is specific to the analog-sensitive TTBK2. Interestingly, both 3-IB-PP1 (10 and 25 μM) and 3-MB-PP1 (25 μM) treatment led to a reduced cilia length compared to DMSO-treated TTBK2 M94G condition (Figure 2C-D). A similar trend (reduced cilia length) was detected when comparing TTBK2 M94G and WT cells treated with a respective ATP analog, although the effect did not reach statistical significance. Noteworthy, we observed some off-target effects of 25 μM 3-IB-PP1 on cilia length in the TTBK2 WT condition, which led us to focus on the 3-MB-PP1 compound in the subsequent experiments. To corroborate our findings, we examined CP110 levels following 3-MB-PP1 treatment. While we detected no effect on CP110 mother centriole levels in WT, we observed an increase in CP110 levels in the TTBK2 M94G condition (Suppl. Fig.2). This is in line with the established role of CP110 as a negative regulator of ciliogenesis initiation and the role of TTBK2 in promoting its removal from the mother centriole distal end in a kinase activity-dependent manner (Goetz et al. 2012).

**Figure 2:**
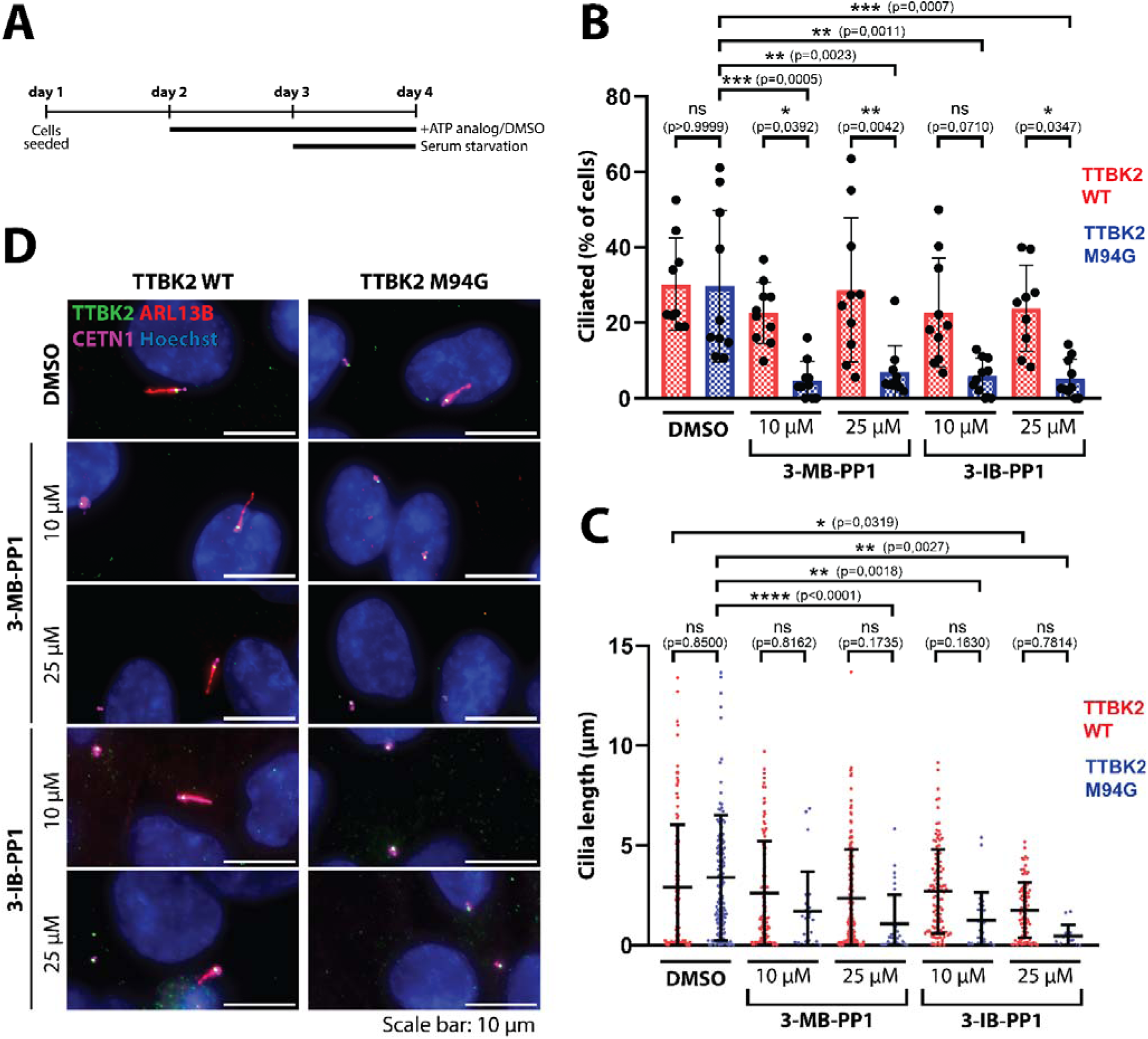
**(A)** Design of the experiment done in TTBK2 WT and TTBK2 M94G hTERT-RPE1 cells, analyzed using IF microscopy. (**B**) Quantification of the percentage of ciliated cells (visualized with ARL13B antibody staining). (**C**) Analysis of the length of ARL13B-positive cilia in the indicated conditions. (**D**) Representative IF microscopy images showing ARL13B (red) positive cilia. CETN1 (purple) was used to visualize centrioles; Hoechst nuclear staining is in blue. Note that both TTBK2 (shown in green) WT and M94G readily localized to the mother centriole across all conditions. Each graph represents a summary of 3 independent experiments and depicts data as mean ±SD. Individual datapoints in **B** represent percentage of ciliated cells per field of view, while in **C** each datapoint represents an individual cilium. Results of statistical analysis (one-way ANOVA with Tukey’s multiple comparisons test) are presented as P-values (*P<0.05, **P<0.01, ***P<0.001, ****P<0.0001).

In sum, our results established that treatment with ATP analogs 3-MB-PP1 or 3-IB-PP1 impairs primary cilia formation by specifically targeting the activity of TTBK2 M94G. In addition, 3-MB-PP1 also showed TTBK2 M94G-specific effects on primary cilia length and CP110 levels. To the best of our knowledge, the analog-sensitive variant of TTBK2 reported here represents the first example of a cilia-related kinase engineered to respond to pharmacological inhibition by bulky ATP analogs.

### ATP analog system reveals rapid dynamics of TTBK2-regulated events

One of the outstanding questions related to TTBK2 biology is the timing and dynamics of TTBK2-regulated events. To probe the potential of the TTBK2 analog-sensitive variant to bring new insight into that, we carried out ATP analog treatment and wash-out experiments, with the overall idea to use the TTBK2 inhibition as “synchronization point”, by halting the ciliogenesis at its very early stage and subsequently analyzing the individual events and their dynamics following induced restart of ciliogenesis (Figure 3A).

**Figure 3:**
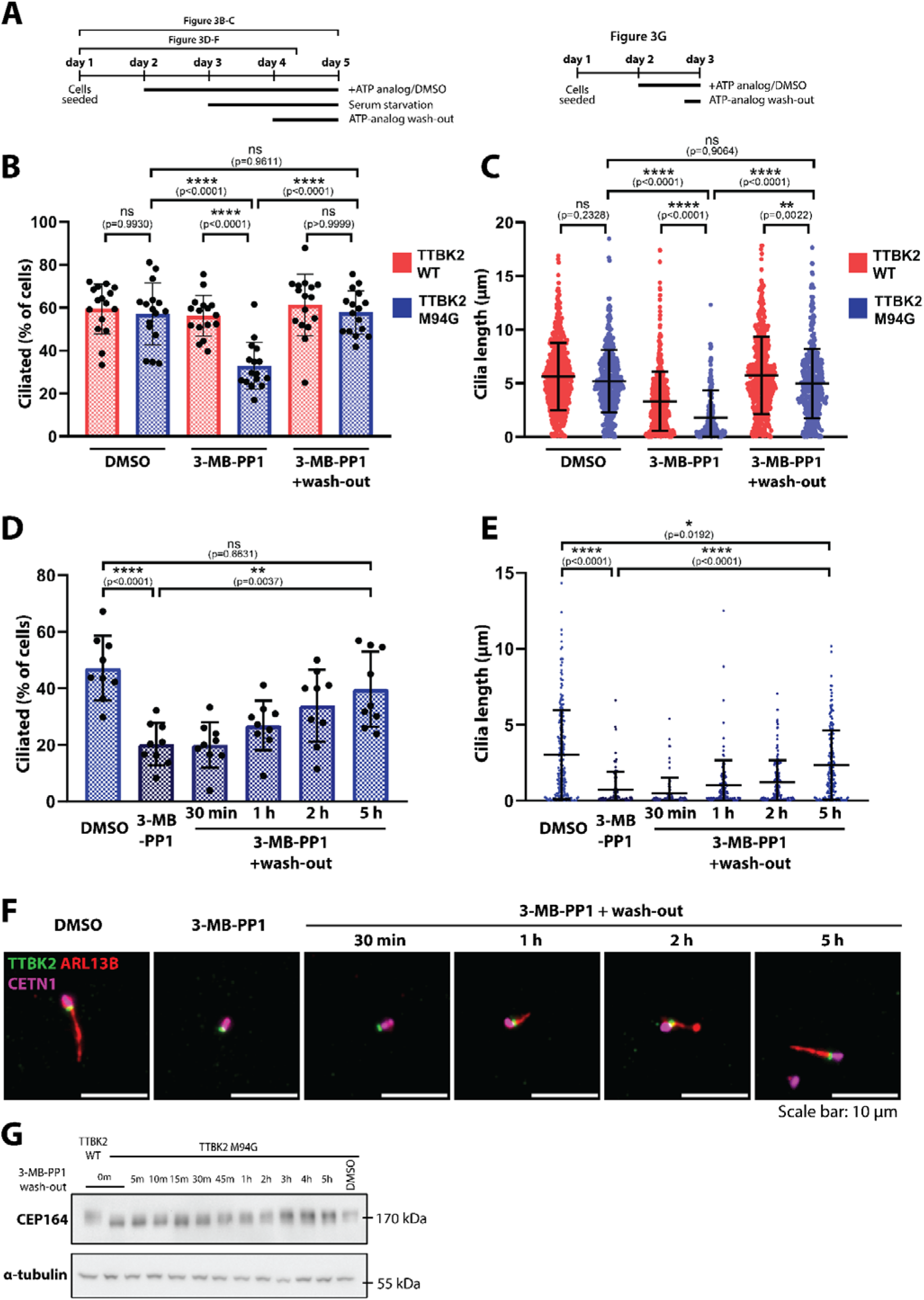
**(A)** Design of the presented experiments done in TTBK2 WT and TTBK2 M94G hTERT-RPE1. Outline of the IF microscopy-based experiment in Fig. 3B-F is on the left; design of the WB experiment in Fig. 3G is on the right. 3-MB-PP1 was used at a 25 μM concentration. (**B**) Quantification of the percentage of ciliated cells (visualized using ARL13B antibody) and (**C**) primary cilia length following 24 hours of 3-MB-PP1 wash-out. (**D**) Quantification of the percentage of ciliated cells and (**E**) cilia length within the first hours upon wash-out of 3-MB-PP1 in TTBK2 M94G hTERT-RPE1 cells. Representative IF microscopy images are shown in (**F**) ARL13B (red) staining was used to detect cilia, CETN1(purple) to visualize centrioles. TTBK2 signal is shown in green; Hoechst nuclear staining is in blue. (**G**) Western blot analysis of CEP164 phosphorylation-dependent mobility shift, representative blot from 2 individual experiments. Please note the gradual increase of phosphorylated CEP164 within the first hours after 3-MB-PP1 wash-out. Alpha-tubulin detection was used as a loading control. Each graph represents a summary of 3 independent experiments and depicts data as mean ±SD. Individual datapoints in **B**,**D** represent the percentage of ciliated cells per field of view, while in **C**,**E** each datapoint represents an individual cilium. Results of statistical analysis (one-way ANOVA with Tukey’s multiple comparisons test) are presented as P-values (*P<0.05, **P<0.01, ***P<0.001, ****P<0.0001).

First, to establish that inhibition of TTBK2 M94G by 3-MB-PP1 is reversible, we treated the cells with 3-MB-PP1 (10 or 25 μM) or DMSO from day 2, followed by serum starvation from day 3 and eventual wash-out of 3-MB-PP1 in selected conditions for the last 24 h (Figure 3A). Importantly, we observed that while the percentage of ciliated cells was notably reduced in 3-MB-PP1-treated M94G condition, it was fully restored to control levels (3-MB-PP1 treated TTBK2 WT or DMSO treated M94G) following the ATP analog wash-out (Figure 3B, Suppl. Fig.3B). Similarly, wash-out of 3-MB-PP1 significantly restored primary cilia length in M94G condition (Figure 3C, Suppl. Fig.3B), confirming the inhibitory effects of 3-MB-PP1 on the analog-sensitive TTBK2 and ciliogenesis were indeed reversible. An experiment with 10 μM 3-MB-PP1 treatment led to similar, yet less pronounced, effects (Suppl. Fig.3C-D), and we therefore used 3-MB-PP1 in 25 μM concentration in all further experiments unless stated otherwise. Of note, in both cases we observed some effect on cilia length even in the 3-MB-PP1-treated WT cells, indicating the appearance of off-target activities during the prolonged treatment.

Having established that 24 hours after the reactivation of TTBK2 by ATP-analog wash-out is sufficient to restore the rate of ciliated cells to control levels, we then focused on the analysis of primary cilia assembly within the first hours after the wash-out of 25 μM 3-MB-PP1 (Figure 3A). Interestingly, additional ARL13B positive cilia started to appear 1-2 hours after the 3-MB-PP1 wash-out, and within 5 hours, the percentage of ciliated cells reached the level of DMSO-treated cells (Figure 3D and 3F). Similarly, the length of cilia was gradually increasing to near control levels during the 5-hour window (Figure 3E and 3F). These data indicate that the timing of cilia assembly following TTBK2 inhibition and wash-out is notably faster than the assembly times reported for primary cilia in asynchronous (Ford et al. 2018; Westlake et al. 2011; Kim et al. 2015; Cao et al. 2012) or synchronized cells (Azizzanjani et al. 2025). For example, serum starvation was shown to trigger axoneme formation within 12–24 hours (Westlake et al. 2011), or within 10-12 hours in cells synchronized in metaphase prior to the starvation (Azizzanjani et al. 2025). Faster times for cilium assembly (6-13 hours) were reported for cytochalasin D-mediated destabilization of actin filaments (Kim et al. 2015) or for cilium assembly following G1 cell cycle phase entry (Ford et al. 2018).

To corroborate our findings regarding the dynamics of cilium assembly following ATP analog wash-out, we decided to evaluate the phosphorylation status of CEP164, an interacting partner and substrate of TTBK2 (Čajánek and Nigg 2014). Similarly to DVL2, phosphorylation of CEP164 can be monitored by its reduced mobility on SDS-PAGE (Bernatik et al. 2020). Indeed, treatment with 3-MB-PP1 (Suppl. Fig. 3A) led to a mobility shift of CEP164 in TTBK2 M94G but not WT conditions already after 1-2 hours (Suppl. Fig. 3E). Similarly, following the wash-out of 3-MB-PP1 (Figure 3A), we observed a gradual increase in CEP164 phosphorylation levels, with the mobility pattern of CEP164 being indistinguishable from control cells after 3 hours (Figure 3G). These data provide biochemical evidence of TTBK2 reactivation following ATP analog washout and align well with the observed rapid induction of primary cilium assembly.

In sum, our model system based on TTBK2 M94G inhibition and reactivation represents a tool for inducing rapid primary cilium assembly. We cannot formally exclude that residual activity of TTBK2 M94G in the presence 3-MB-PP1 allows some initial events of ciliogenesis to take place, which may contribute to the observed rapid dynamics of cilium re-assembly.

### TTBK2 inhibition leads to rapid loss of IFT88 and cilia shortening

Recent work has linked TTBK2 activity to the regulation of primary cilia maintenance (Lin et al. 2024; Nguyen and Goetz 2022; Benk Vysloužil et al. 2025). To test whether the 3-MB-PP1-mediated inhibition of TTBK2 M94G is sufficient to affect cilia maintenance, we performed an experiment where cells were deprived of serum 24 hours prior to the 3-MB-PP1 treatment (Suppl. Fig.4A). Indeed, we observed a drop in the percentage of ciliated cells in 3-MB-PP1 treated TTBK2 M94G over TTBK2 WT or DMSO-treated controls (Suppl. Fig. 4B,D). However, as we found no difference between untreated TTBK2 M94G cells fixed on day 4 and 3-MB-PP1 treated TTBK2 M94G cells fixed on day 5, it is difficult to unambiguously relate the difference between M94G and WT conditions to cilia maintenance defect. On the other hand, we found that the 3-MB-PP1 treatment impaired the length of assembled cilia in the M94G condition, in line with our previous observations (Suppl. Fig.4C-D), suggesting an effect of M94G inhibition on cilia maintenance in the end.

To corroborate the effects of 3-MB-PP1 mediated inhibition of TTBK2 M94G on the maintenance of cilia, we adjusted our protocol to assess a more rapid response (Fig. 4A). Intriguingly, we detected a reduction in the cilia length of 3-MB-PP1 treated TTBK2 M94G, but not WT control, as early as 2 hours after 3-MB-PP1 treatment (Fig.4B-C, Suppl. Fig.4E-F). Interestingly, we detected an even more rapid response to 3-MB-PP1 mediated TTBK2 M94G inhibition when analyzing levels of IFT88, a component of the IFT B complex essential for ciliogenesis (Pazour et al. 2000). As shown in Figure 4D-E, IFT88 levels at the mother centriole dropped rapidly, with a significant reduction already evident after 30 minutes of treatment. The difference from the control further increased after 2 hours. Importantly, the level of IFT88 at the mother centriole remained unaffected in 3-MB-PP1-treated TTBK2 WT cells (Suppl. Fig.4G-H).

**Figure 4:**
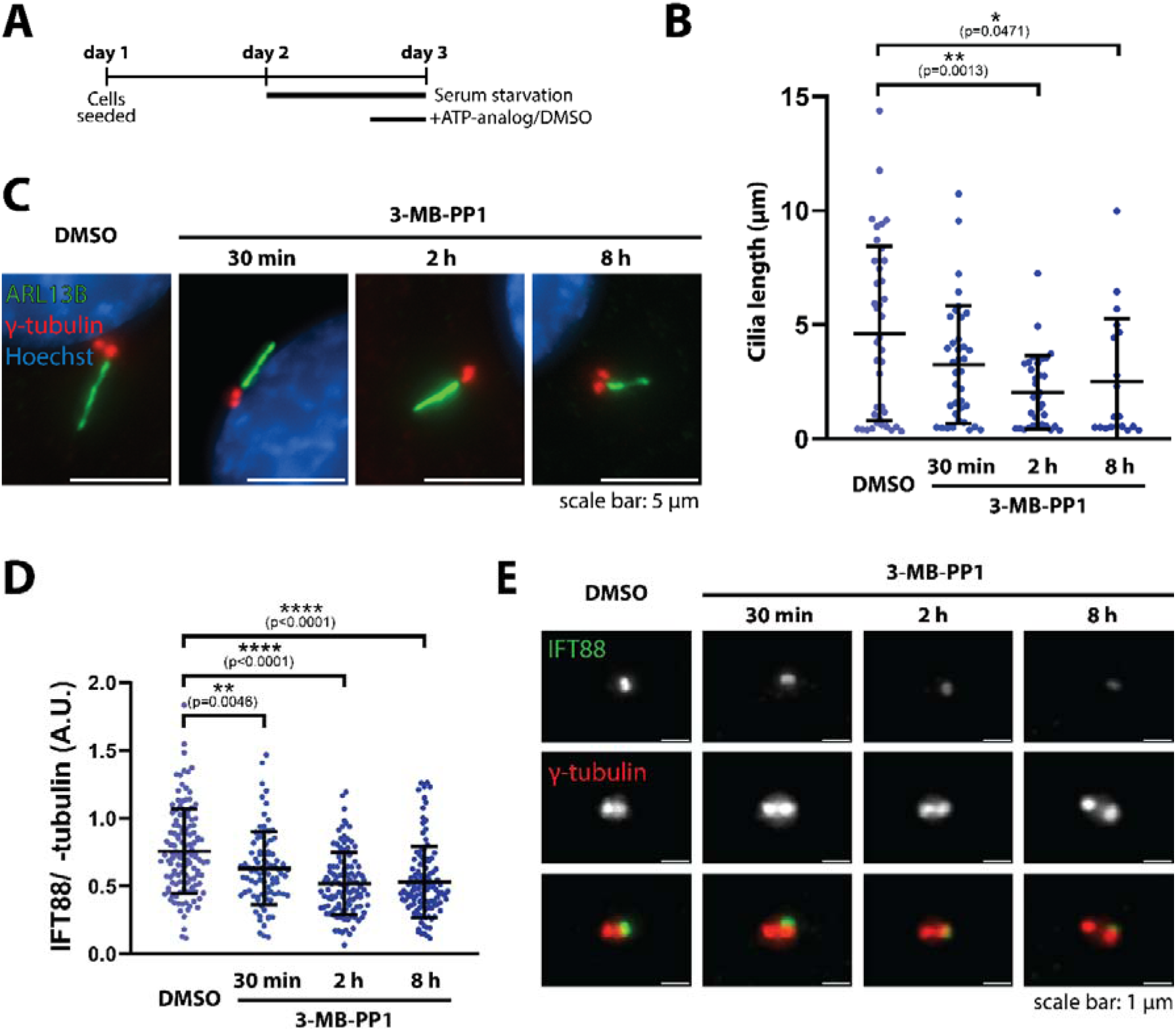
**(A)** Design of the presented experiments done in TTBK2 M94G hTERT-RPE1. 3-MB-PP1 was used at a 25 μM concentration. (**B**) Quantification of the length of primary cilia (visualized using ARL13B antibody) after 3-MB-PP1 treatment. (**C**) Representative IF microscopy images of cilia (detected using ARL13B antibody staining, green) in the indicated conditions. Gamma-tubulin staining (red) visualized centrosomes; Hoechst nuclear staining is shown in blue. (**D**) Quantification of relative IFT88 levels (normalized to the intensity of γ-tubulin) at the mother centriole upon 3-MB-PP1 treatment. Representative IF microscopy images are shown in (**E**). IFT88 staining is shown in green, γ-tubulin staining is in red. Each graph represents a summary of 3 independent experiments and depicts data as mean ±SD. Individual datapoints in **B** represent the percentage of ciliated cells per field of view, while in **D** each datapoint represents an individual cilium. Results of statistical analysis (one-way ANOVA with Tukey’s multiple comparisons test) are presented as P-values (*P<0.05, **P<0.01, ***P<0.001, ****P<0.0001).

While these data are correlative, they support a model in which impaired IFT recruitment to the cilia base leads to rapid cilia shortening and eventual disassembly in vertebrate cells, as reported for NIH-3T3 cells following acute inhibition of IFT motor, heterotrimeric kinesin II (Engelke et al. 2019). In sum, our observations further expand the potential of the engineered ATP-analog-sensitive TTBK2 variant for investigating mechanisms governing the maintenance of primary cilia.

It should be noted that the analog-sensitive TTBK2-based model system has shortcomings, too. Perhaps the most relevant limitation is off-target activity, manifesting in some of our long-term experiments as an effect of the used ATP analog on primary cilia in the TTBK2 WT condition (Fig.3C, Suppl. Fig.4C). This observation confirms and extends a recent report describing off-target effects of PP1-derived ATP analogs (Gicquel et al. 2026) and reinforces the need for careful experiment planning and the use of appropriate controls. Importantly, as kinase inhibitors in general are infamous for off-target effects when used in high concentrations (Bain et al. 2007), the ATP analog-sensitive kinase model offers the ultimate specificity control in the WT kinase treated with the same compound.

## Acknowledgements

We thank David Benk Vysloužil for the initial work on this project and Vítězslav Bryja for sharing reagents with us. We acknowledge the core facility CELLIM supported by the Czech-BioImaging large RI project (LM2023050 funded by MEYS CR) for their support with obtaining scientific data presented in this paper.

## Author contributions

AL – designed and performed experiments, analyzed data, and together with LČ wrote the manuscript; LČ – conceived and supervised the project, acquired funding.

## Data Availability Statement

The original data are available upon reasonable request.

## Supplementary material

**Supplementary Figure 1:**
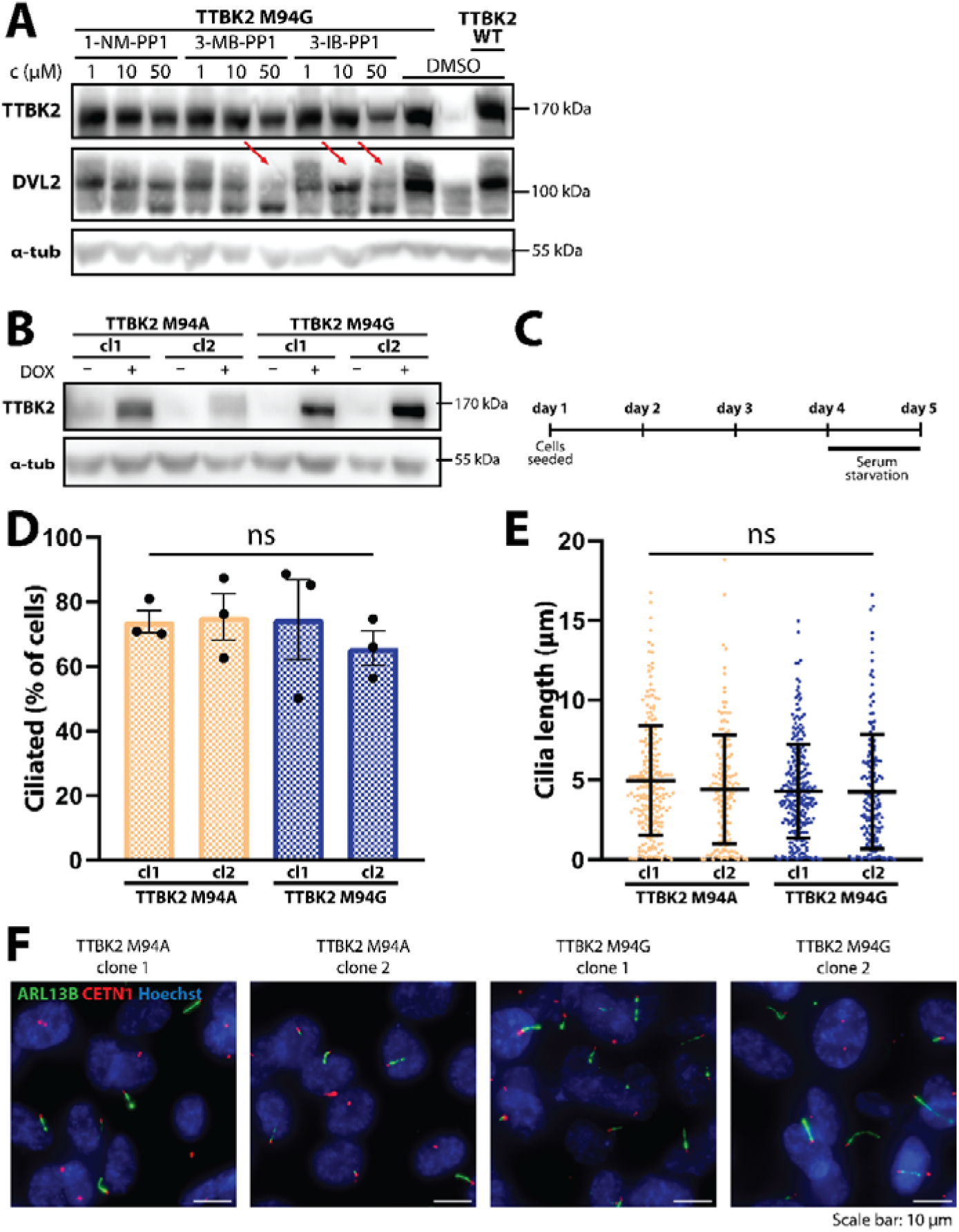
(**A**) Western blot analysis of the potency of available synthetic ATP-analog compounds to block catalytic activity of TTBK2 M94G. TTBK2 KO HEK293 cells transfected with Flag-tagged TTBK2 (either WT or M94G) together with TTBK2’s substrate DVL2 were treated with ATP-analog compounds as indicated. Please note the reduction of upshifted DVL2 band upon treatment with 3-MB-PP1 (50 μM) and 3-IB-PP1 (10 μM and 50 μM) (red arrows). Alpha-tubulin detection was used as a loading control. Representative blot from 2 individual experiments. (**B**) Western blot validation of DOX-inducible TTBK2 expression in M94A and TTBK2 M94G hTERT-RPE1 stable cell lines (two clones per stable line examined). Alpha-tubulin detection was used as a loading control. (**C**) Design of the presented experiments done in TTBK2 M94A and TTBK2 M94G hTERT-RPE1 cells. (**D**) Percentage of ciliated cells and (**E**) length of cilia in the generated cell lines. Representative IF microscopy images are shown in (**F**). Cilia are marked by ARL13B staining (green), centrioles are visualized by CETN1 staining (red); Hoechst nuclear staining is shown in blue. Each graph represents one experiment and depicts data as mean ±SD. Individual datapoints in **D** represent percentage of ciliated cells per field of view, while in **E** each datapoint represents an individual cilium. Statistical analysis was performed using one-way ANOVA with Tukey’s multiple comparisons test. P=0.05 was set as the threshold for statistical significance.

**Supplementary Figure 2:**
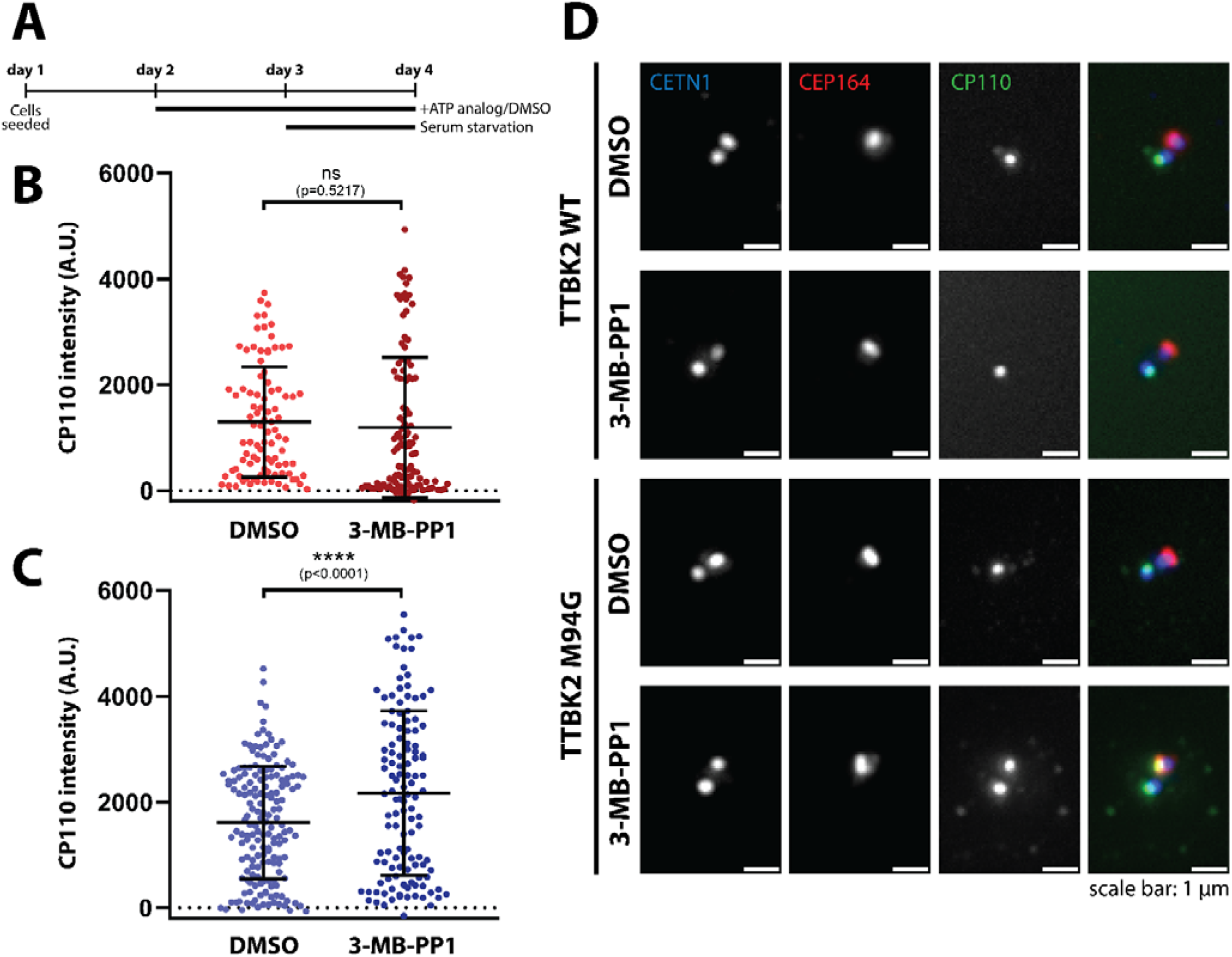
(**A**) Design of the presented experiments done in TTBK2 WT and TTBK2 M94G hTERT-RPE1. 3-MB-PP1 was used at a 25 μM concentration. Quantification of CP110 levels at the mother centriole upon 3-MB-PP1 treatment of (**B**) TTBK2 WT and (**C**) TTBK2 M94G hTERT-RPE1 cells. Representative IF microscopy images are shown in (**D**). Centrioles are visualized with CETN1 (blue), mother centriole is marked with CEP164 (red). CP110 staining is in green. Each graph represents a summary of 2 independent experiments and depicts data as mean ±SD, each datapoint represents one centriole. Results of statistical analysis (one-way ANOVA with Tukey’s multiple comparisons test) are presented as P-values (*P<0.05, **P<0.01, ***P<0.001, ****P<0.0001).

**Supplementary Figure 3:**
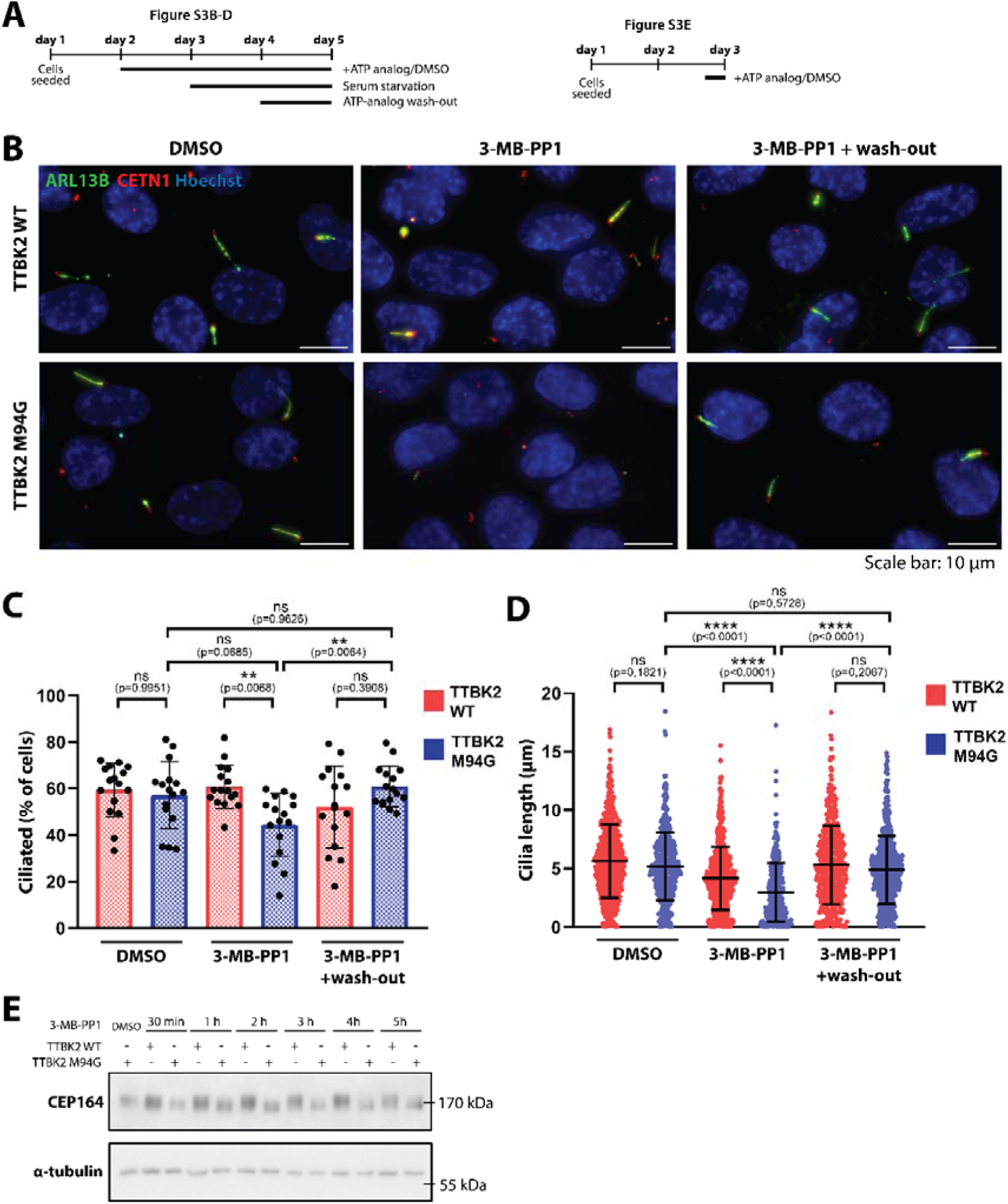
(**A**) Design of the presented experiments done in TTBK2 WT and TTBK2 M94G hTERT-RPE1. 3-MB-PP1 was used in 10 μM (Suppl. Fig.3C-D) or 25 μM (Suppl. Fig.3B,E) concentration. (**B**) Representative IF microscopy images showing ARL13B (green) positive cilia. CETN1 (red) was used to visualize centrioles; Hoechst nuclear staining is in blue. (**C**) Quantification of the percentage of ciliated cells (visualized using ARL13B antibody) and (**D**) primary cilia length following 24 hours of 3-MB-PP1 wash-out. (**E**) Western blot analysis of CEP164 phosphorylation-dependent mobility shift, representative blot from 2 individual experiments. Please note the gradual decrease of phosphorylated CEP164 within the first hours after 3-MB-PP1 wash-out. Alpha-tubulin detection was used as a loading control. Each graph represents a summary of 3 independent experiments and depicts data as mean ±SD. Individual datapoints in **C** represent percentage of ciliated cells per field of view, while in **D** each datapoint represents an individual cilium. Results of statistical analysis (one-way ANOVA with Tukey’s multiple comparisons test) are presented as P-values (*P<0.05, **P<0.01, ***P<0.001, ****P<0.0001).

**Supplementary Figure 4:**
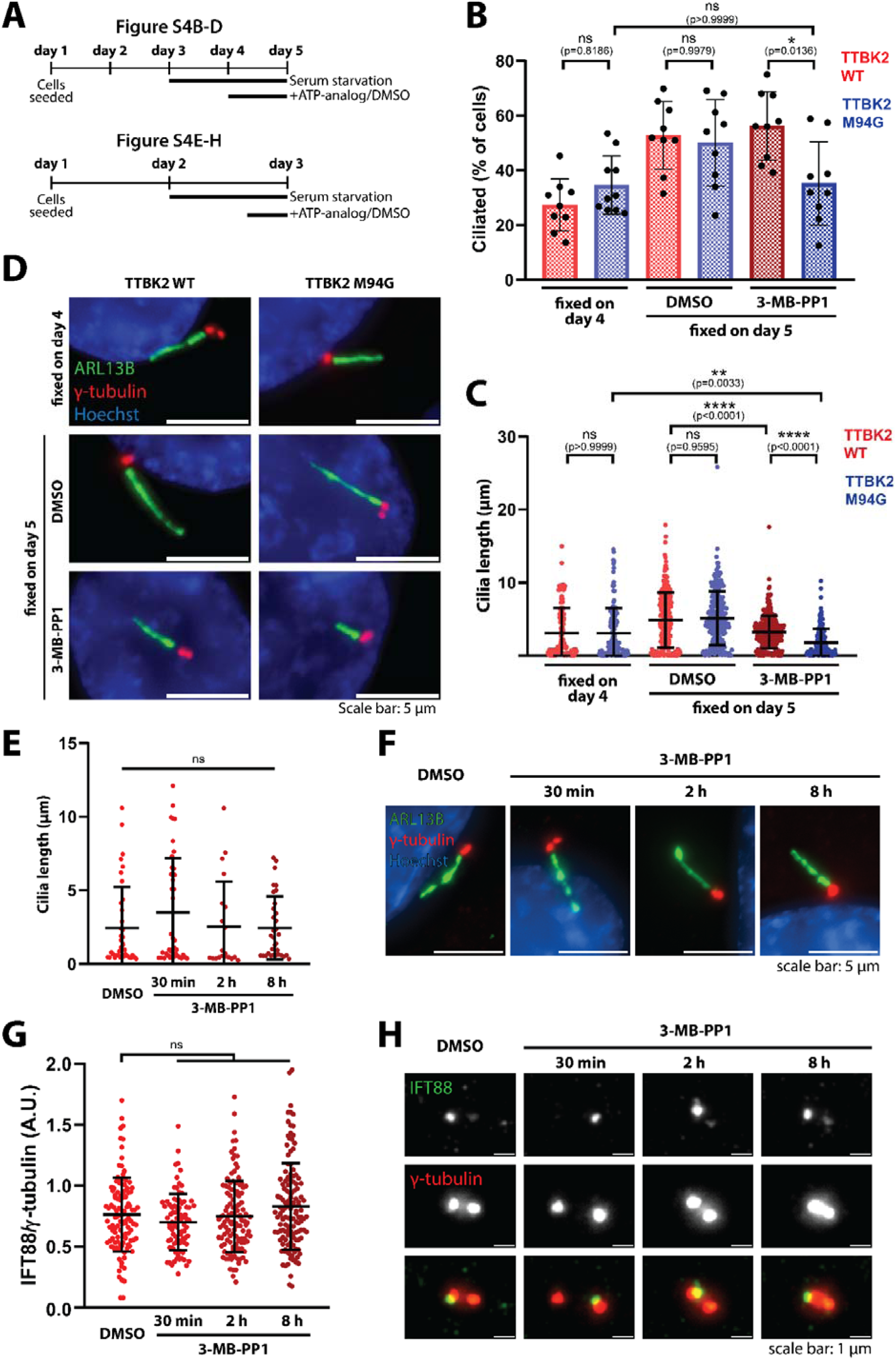
(**A**) Design of the presented experiments done in TTBK2 WT and TTBK2 M94G hTERT-RPE1. 3-MB-PP1 was used at a concentration of 25 μM. (**B**) Quantification of the percentage of ciliated cells (visualized using ARL13B antibody) and (**C**) primary cilia length prior to or following 3-MBPP1 treatment. Cells were either fixed on day 4 (prior to the treatment) or treated with 3-MB-PP1 for 24 hours. Representative IF microscopy images are shown in (**D**). ARL13B (green) staining was used to detect cilia, γ-tubulin (red) to visualize centrioles. Hoechst nuclear staining is in blue. (**E**) Quantification of the length of primary cilia (visualized using ARL13B antibody) after 3-MB-PP1 treatment of TTBK2 WT hTERT-RPE1 cells. (**F**) Representative IF microscopy images of cilia (detected using ARL13B antibody staining, green) in the indicated conditions. Gamma-tubulin staining (red) visualized centrosomes; Hoechst nuclear staining is shown in blue. (**G**) Quantification of relative IFT88 levels (normalized to the intensity of γ-tubulin) at the mother centriole upon 3-MB-PP1 treatment of TTBK2 WT hTERT-RPE1 cells. Representative IF microscopy images are shown in (**H**). IFT88 staining is shown in green, γ-tubulin staining is in red. Each graph represents a summary of 3 independent experiments and depicts data as mean ±SD. Individual datapoints in **B** represent the percentage of ciliated cells per field of view, in **C**,**E** each datapoint represents an individual cilium, and in **G**, each datapoint represents one centriole. Results of statistical analysis (one-way ANOVA with Tukey’s multiple comparisons test) are presented as P-values (*P<0.05, **P<0.01, ***P<0.001, ****P<0.0001).

