## Supplementary figures and images for "An Analog-Sensitive Tau Tubulin Kinase 2 Enables Acute Chemical-Genetic Control of Primary Cilia"

### Supplementary material

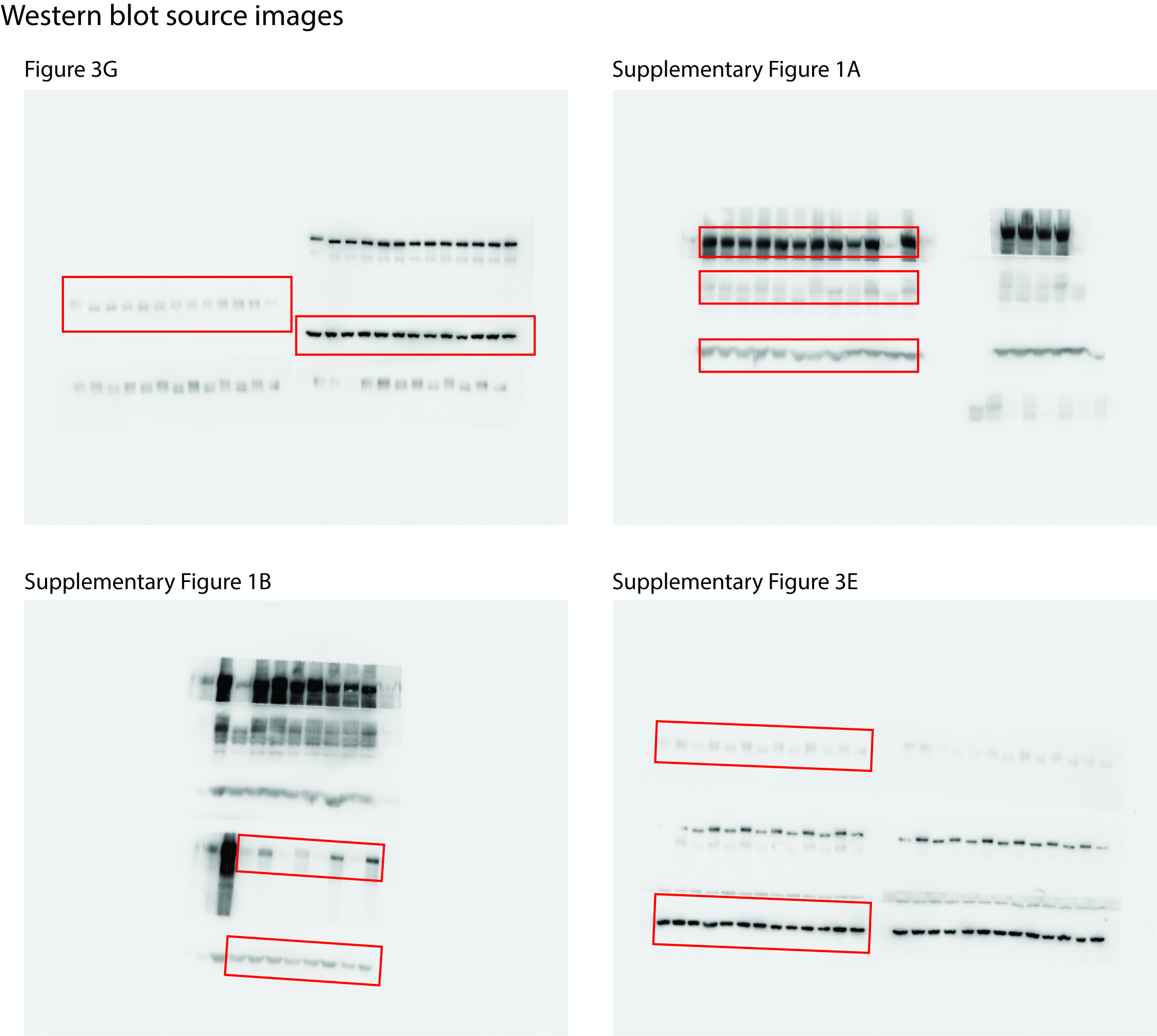
